# Sensitivity of BA.2.86 to prevailing neutralising antibody responses

**DOI:** 10.1101/2023.09.02.556033

**Authors:** Daniel J Sheward, Yiqiu Yang, Michelle Westerberg, Sofia Öling, Sandra Muschiol, Kenta Sato, Thomas P Peacock, Gunilla B Karlsson Hedestam, Jan Albert, Ben Murrell

**Affiliations:** Department of Microbiology, Tumor and Cell Biology, Karolinska Institutet, Stockholm, Sweden; Department of Clinical Microbiology, Karolinska University Hospital, Stockholm, Sweden; Department of Infectious Disease, Imperial College London, London, UK; The Pirbright Institute, Woking, UK, GU24 0NF

## Abstract

A new SARS-CoV-2 variant, designated BA.2.86, has recently emerged with over 30 spike mutations relative to its parental BA.2, raising questions about its degree of resistance to neutralising antibodies. Using a spike-pseudotyped virus model we characterise neutralisation of BA.2.86 by clinically relevant monoclonal antibodies and by two cohorts of serum sampled from Stockholm, including both a recent cohort, and one sampled prior to the arrival of XBB in Sweden.

## Main Text

After a prolonged period of near-complete global dominance of the recombinant XBB family of SARS-CoV-2 lineages, a substantially mutated sublineage of BA.2 has recently emerged. Designated BA.2.86^1^, the geographic origin of this sublineage is currently unclear, and as of Sept 1st, 2023, BA.2.86 sequences have been observed in 9 countries: Israel, Denmark, the USA, the United Kingdom, South Africa, Portugal, Sweden, Canada, and France^2^.

With a long branch of unobserved evolution (Fig. A) including over 30 mutations in the spike protein relative to BA.2, with many at key antigenic sites (Fig. B and table SI1), its emergence is reminiscent of the initial emergence of Omicron/BA.1^3^, raising immediate questions about whether it is sensitive to any previously approved clinical monoclonal antibodies, and the degree to which it is able to escape antibody responses in the current setting where individual exposure histories are a complex combination of multiple immunizations and multiple past SARS-CoV-2 infections.

**Figure.**
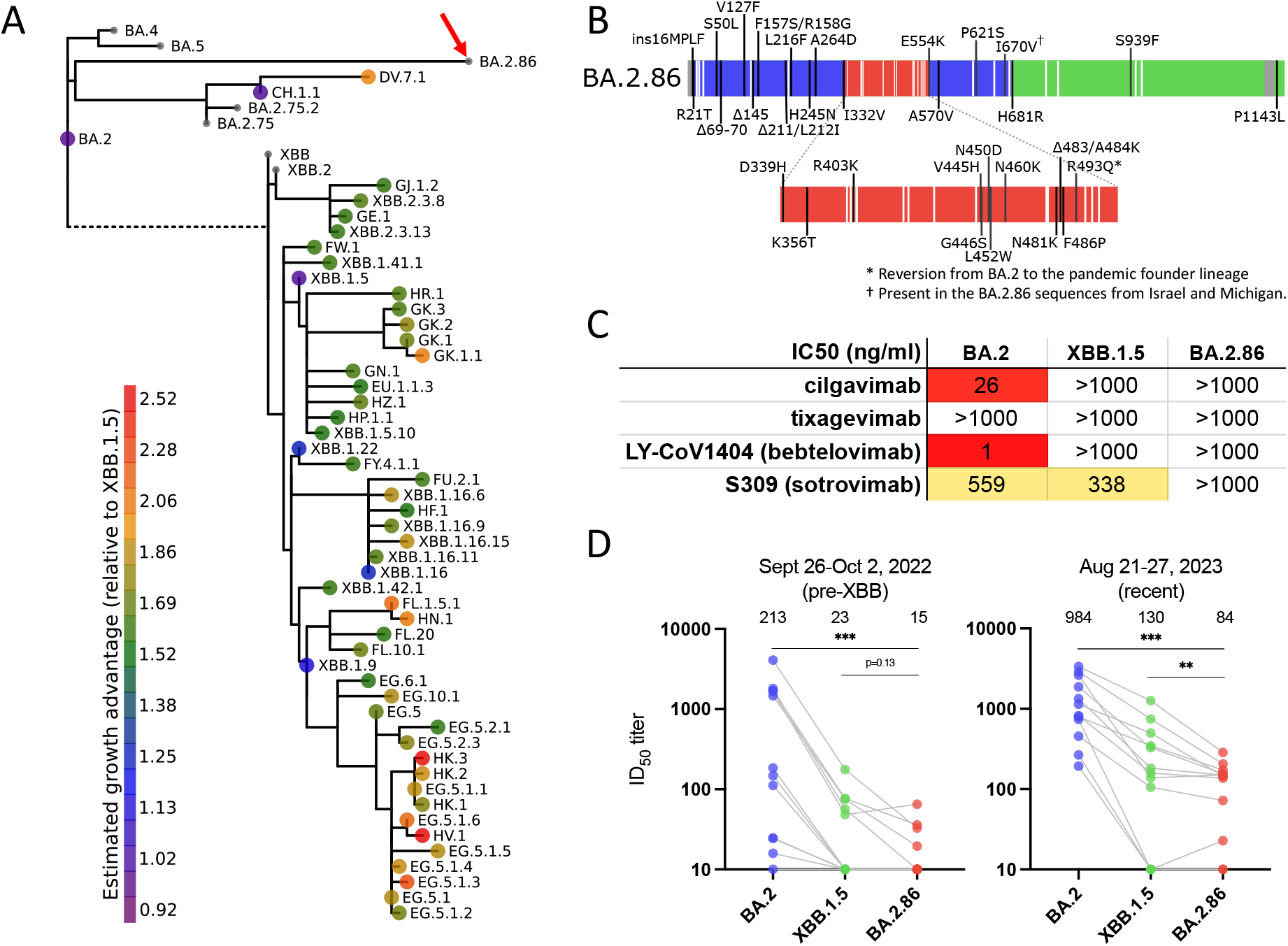
**A**. Phylogeny of currently competing variants, showing the position of BA.2.86 with a red arrow. Circles show estimated variant growth advantage. Small circles denote insufficient data within the previous 100 days to reliably estimate relative growth advantage. **B**. Mutations in BA.2.86 relative to BA.2. **C**. Neutralisation of BA.2, XBB.1.5, and BA.2.86 pseudoviruses by clinically relevant monoclonal antibodies. **D**. Pseudovirus neutralisation by serum samples taken Sept 26 - Oct 2, 2022, prior to the arrival of XBB in Sweden, as well as by current samples (Aug 21-27, 2023).

Using a BA.2.86 spike-pseudotyped virus model (see SI Methods), we find that BA.2.86 is resistant to clinically relevant monoclonal antibodies (Fig. C) tixagevimab and cilgavimab (Evusheld), LY-CoV1404 (bebtelovimab) and S309 (sotrovimab).

To characterise the relative sensitivity of BA.2.86 to current antibody-mediated immunity at the population level, we evaluated BA.2.86 neutralisation by serum sampled from random blood donors in Stockholm during week 34 (Aug. 21 - 27, 2023). Furthermore, to provide insight into the impact of XBB exposure - given that this variant may be included in upcoming, updated vaccines - we also assessed neutralisation of BA.2.86 by serum sampled from blood donors prior to the circulation of XBB (week 39, 2022 - Sep. 26 - Oct. 2). We compare BA.2.86 to the basal BA.2, and the recently dominant XBB.1.5.

Serum sampled ‘pre-XBB’ had substantially lower (4-6-fold) neutralising antibody titers than recently sampled sera across all three variants, with 9 of 12 pre-XBB samples having ID50 titers against BA.2.86 below the limit of detection (20) (Fig D). Across both time points, sera had moderately lower geometric mean neutralising titers against BA.2.86 than against XBB.1.5. Titers to BA.2.86 were also more than 10-fold lower than against its parent, BA.2. Current sera from Sweden had elevated titers against all variants, likely due to prevalent infections with XBB and its descendants. Notably, 8 of 12 recent serum samples exhibited titers over 100 against BA.2.86.

Nothing is yet known about the severity of BA.2.86 and, with diminished global SARS-CoV-2 sequencing efforts, it is too early to forecast whether BA.2.86, or one of its descendants, will outcompete currently circulating lineages. While the occurrence of another Omicron-like emergence event should caution against reducing our surveillance capacity, immune escape of BA.2.86 does not appear to be as extensive as was the case when Omicron emerged in the background of Delta^4^, likely attributable to multiple immunizations and prior infections since.

## Acknowledgements

pCMV-dR8.2 dvpr was a gift from Bob Weinberg (Addgene plasmid # 8455; http://n2t.net/addgene:8455; RRID:Addgene_8455). pBOBI-FLuc was a gift from David Nemazee (Addgene plasmid # 170674; http://n2t.net/addgene:170674; RRID:Addgene_170674).

We thank Roy A Ehling and Sai T Reddy for providing plasmids for antibodies S309 and LY-CoV1404, and Niklas K Björkström for providing tixagevimab and cilgavimab.

We gratefully acknowledge all data contributors, i.e. the Authors and their Originating Laboratories responsible for obtaining the specimens, and their Submitting Laboratories that generated the genetic sequence and metadata and shared via the GISAID Initiative the data on which part of this research is based.

## Supplementary Material

**Table SI 1.**
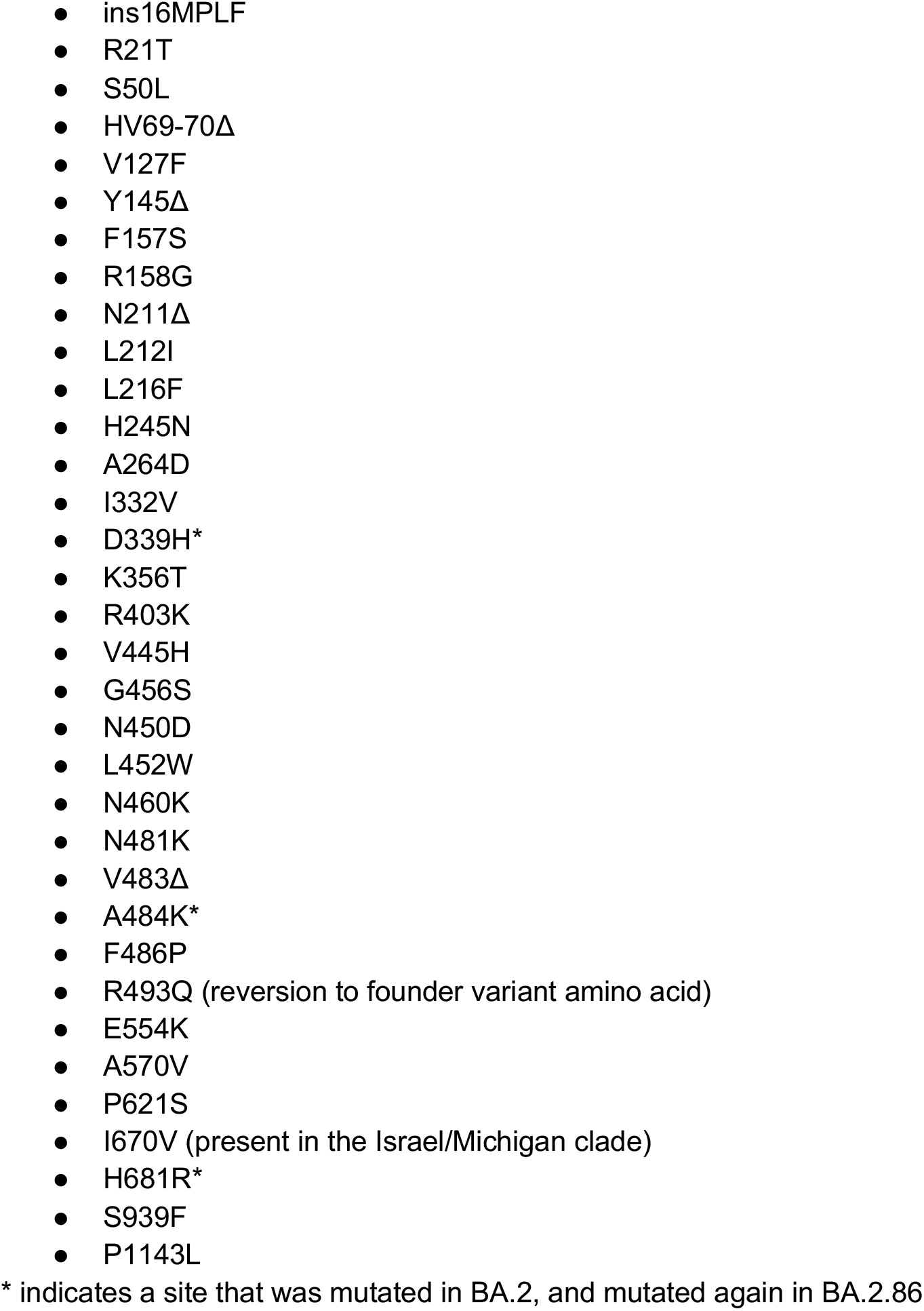
BA.2.86 mutations relative to BA.2:

Nucleotide sequence for codon-optimised BA.2.86 spike construct:

~~~
ATGTTCGTGTTTCTGGTGCTGCTGCCTCTGGTGTCCAGCCAGTGTGTGATGCCACTGTTTAACCTGATCACCACTACACAGTC
GTACACCAACAGCTTTACCAGAGGCGTGTACTACCCCGACAAGGTGTTCAGATCCAGCGTGCTGCACCTGACCCAGGACCTGT
TCCTGCCTTTCTTCAGCAACGTGACCTGGTTCCACGCCATCTCCGGCACCAATGGCACCAAGAGATTCGACAACCCCGTTCTG
CCCTTCAACGACGGGGTGTACTTTGCCAGCACCGAGAAGTCCAACATCATCAGAGGCTGGATCTTCGGCACCACACTGGACAG
CAAGACCCAGAGCCTGCTGATCGTGAACAACGCCACCAACGTGTTTATCAAAGTGTGCGAGTTCCAGTTCTGCAACGACCCCT
TCCTGGACGTCTACCACAAGAACAACAAGAGCTGGATGGAAAGCGAGTCAGGCGTGTACAGCAGCGCCAACAACTGCACCTTC
GAGTACGTGTCCCAGCCTTTCCTGATGGACCTGGAAGGCAAGCAGGGCAACTTCAAGAACCTGCGCGAGTTCGTGTTCAAGAA
CATCGACGGCTACTTCAAGATCTACAGCAAGCACACCCCTATCATCGGCCGGGATTTTCCTCAGGGCTTCTCTGCTCTGGAAC
CCCTGGTGGATCTGCCCATCGGCATCAACATCACCCGGTTTCAGACACTGCTGGCCCTGAATAGAAGCTACCTGACACCTGGT
GATAGCAGCAGCGGATGGACAGCTGGTGCCGCCGATTACTATGTGGGCTACCTGCAGCCTAGAACCTTCCTGCTGAAGTACAA
CGAGAACGGCACCATCACCGACGCCGTGGATTGTGCCCTTGATCCTCTGAGCGAGACAAAGTGCACCCTGAAGTCCTTCACCG
TGGAAAAGGGCATCTACCAGACCAGCAACTTCCGGGTGCAGCCCACCGAATCCATCGTGCGGTTCCCCAATGTGACCAATCTG
TGCCCCTTCCACGAGGTGTTCAATGCCACCAGGTTCGCCTCTGTGTACGCCTGGAACCGGACTCGGATCAGCAATTGCGTGGC
CGACTACTCCGTGCTGTACAACTTCGCCCCCTTCTTCGCCTTCAAGTGCTACGGCGTGTCCCCTACCAAGCTGAACGACCTGT
GCTTCACAAACGTGTACGCCGACAGCTTCGTGATCAAAGGAAATGAAGTGAGTCAGATTGCCCCTGGACAGACAGGCAACATC
GCCGACTACAACTACAAGCTGCCCGACGACTTCACCGGCTGTGTGATTGCCTGGAACAGCAACAAGCTGGACTCCAAACATAG
CGGCAACTACGATTACTGGTACCGGCTGTTCCGGAAGTCCAAGCTGAAGCCCTTCGAGCGGGACATCTCCACCGAGATCTATC
AGGCCGGCAACAAGCCTTGTAAAGGCAAGGGCCCCAACTGCTACTTTCCACTGCAGTCCTACGGCTTTCGGCCCACATATGGC
GTGGGCCATCAGCCCTACAGAGTGGTGGTGCTGAGCTTCGAACTGCTGCATGCCCCTGCCACAGTGTGCGGCCCTAAGAAAAG
CACCAATCTCGTGAAGAACAAATGCGTGAACTTCAACTTCAACGGCCTGACCGGCACAGGCGTGCTGACAAAAAGCAACAAGA
AGTTCCTGCCATTCCAGCAGTTTGGCCGGGATATCGTCGATACCACAGACGCCGTTAGAGATCCCCAGACACTGGAAATCCTG
GACATCACCCCTTGCAGCTTCGGCGGAGTGTCTGTGATCACCCCTGGCACCAACACCAGCAATCAGGTGGCAGTGCTGTACCA
GGGCGTGAACTGTACCGAAGTGAGCGTGGCCATTCACGCCGATCAGCTGACACCTACATGGCGGGTGTACTCCACCGGCAGCA
ATGTGTTTCAGACCAGAGCCGGCTGTCTGATCGGAGCCGAGTACGTGAACAATAGCTACGAGTGCGACATCCCCATCGGCGCT
GGCGTGTGTGCCAGCTACCAGACACAGACAAAGAGCCGCAGACGGGCCAGATCTGTGGCCAGCCAGAGCATCATTGCCTACAC
AATGTCTCTGGGCGCCGAGAACAGCGTGGCCTACTCCAACAACTCTATCGCTATCCCCACCAACTTCACCATCAGCGTGACCA
CAGAGATCCTGCCTGTGTCCATGACCAAGACCAGCGTGGACTGCACCATGTACATCTGCGGCGATTCCACCGAGTGCTCCAAC
CTGCTGCTGCAGTACGGCAGCTTCTGCACCCAGCTGAAAAGAGCCCTGACAGGGATCGCCGTGGAACAGGACAAGAACACCCA
AGAGGTGTTCGCCCAAGTGAAGCAGATCTACAAGACCCCTCCTATCAAGTACTTCGGCGGCTTCAATTTCAGCCAGATTCTGC
CCGATCCTAGCAAGCCCAGCAAGCGGAGCTTCATCGAGGACCTGCTGTTCAACAAAGTGACACTGGCCGACGCCGGCTTCATC
AAGCAGTATGGCGATTGTCTGGGCGACATTGCCGCCAGGGATCTGATTTGCGCCCAGAAGTTTAACGGACTGACAGTGCTGCC
TCCTCTGCTGACCGATGAGATGATAGCCCAGTACACATCTGCCCTGCTGGCCGGCACAATCACAAGCGGCTGGACATTTGGAG
CTGGCGCCGCTCTGCAGATCCCCTTTGCTATGCAGATGGCCTACAGATTCAACGGCATCGGAGTGACCCAGAATGTGCTGTAC
GAGAACCAGAAGCTGATCGCCAACCAGTTCAACAGCGCCATCGGCAAGATCCAGGACAGCCTGTTCAGCACAGCAAGCGCCCT
GGGAAAGCTGCAGGACGTGGTCAACCACAATGCCCAGGCACTGAACACCCTGGTCAAGCAGCTGTCCTCCAAGTTCGGCGCCA
TCAGCTCTGTGCTGAACGATATCCTGAGCAGACTGGACAAGGTGGAGGCCGAGGTGCAGATCGACAGACTGATCACAGGCAGA
CTGCAGAGCCTCCAGACATACGTGACCCAGCAGCTGATCAGAGCCGCCGAGATTAGAGCCTCTGCCCATCTGGCCGCCACCAA
GATGTCTGAGTGTGTGCTGGGCCAGAGCAAGAGAGTGGACTTTTGCGGCAAGGGCTACCACCTGATGAGCTTCCCTCAGTCTG
CCCCTCACGGCGTGGTGTTTCTGCACGTGACATACGTTCCCGCTCAAGAGAAGAATTTCACCACCGCTCCAGCCATCTGCCAC
GACGGCAAAGCCCACTTTCCTAGAGAAGGCGTGTTCGTGTCCAACGGCACCCATTGGTTCGTGACACAGCGGAACTTCTACGA
GCCCCAGATCATCACCACCGACAACACCTTCGTGTCTGGCAACTGCGACGTCGTGATCGGCATTGTGAACAATACCGTGTACG
ACCCTCTGCAGTTGGAGCTGGACAGCTTCAAAGAGGAACTGGACAAGTACTTTAAGAACCACACAAGCCCCGACGTGGACCTG
GGCGATATCAGCGGAATCAATGCCAGCGTCGTGAACATCCAGAAAGAGATCGACCGGCTGAACGAGGTGGCCAAGAATCTGAA
CGAGAGCCTGATCGACCTGCAAGAACTGGGGAAGTACGAGCAGTACATCAAGTGGCCCTGGTACATCTGGCTGGGCTTTATCG
CCGGACTGATTGCCATCGTGATGGTCACAATCATGCTGTGTTGCATGACCAGCTGCTGTAGCTGCCTGAAGGGCTGTTGTAGC
TGTGGCAGCTGCTGCTGA
~~~

## Methods

### Cell culture

HEK293T cells (ATCC CRL-3216) and HEK293T-ACE2 cells (stably expressing human ACE2) were cultured in Dulbecco’s Modified Eagle Medium (high glucose, with sodium pyruvate) that was supplemented with 10% fetal bovine serum, 100 units/ml Penicillin, and 100 μg/ml Streptomycin. Cultures were maintained in a humidified 37°C incubator (5% CO2).

### SARS-CoV-2 spike constructs

A codon-optimised BA.2.86 spike was assembled from overlapping fragments (IDT) by Gibson Assembly^1^ and confirmed by Sanger sequencing. BA.2.86 AA mutations, relative to BA.2, are shown in table SI 1. The BA.2.86 pseudovirus construct used here includes those BA.2.86 mutations (including I670V, present in the Israel/Michigan clade), a 19AA tail truncation (to promote spike incorporation), and an incidental mutation N1023H, which is distal to any known antigenic sites, not accessible to antibodies in the pre-fusion spike conformation, and does not disrupt entry. The XBB.1.5 spike-encoding plasmid was generated by multi-site directed mutagenesis as previously described^2^ and subsequently confirmed by Sanger sequencing.

### Pseudovirus Neutralisation Assay

Pseudovirus neutralisation assays were performed as previously^2^. Briefly, spike-pseudotyped lentivirus particles were generated by the co-transfection of HEK293T cells with each spike plasmid, together with an HIV gag-pol packaging plasmid (Addgene #8455), and a transfer plasmid encoding firefly luciferase (Addgene #170674) using polyethylenimine. Neutralisation was assessed in HEK293T-ACE2 cells. Pseudoviruses titrated to produce approximately 100,000 RLU were incubated with serial 3-fold dilutions of serum or monoclonal antibody for 60 minutes at 37°C in black-walled 96-well plates. 10,000 HEK293T-ACE2 cells were then added to each well, and plates were incubated at 37°C for approximately 48 hours. Samples were tested against all variants ‘head-to-head’ using the same dilutions. Luminescence was measured on a GloMax Navigator Luminometer (Promega) using Bright-Glo luciferase substrate (Promega). Neutralisation was calculated relative to the average of 8 control wells infected in the absence of antibody. ID50 values were calculated in Prism v9 (GraphPad Software) by fitting a four-parameter logistic curve.

### Monoclonal antibodies

Cilgavimab and tixagevimab were evaluated as their clinical formulations. LyCoV1404 and S309 were expressed and purified as previously described^3^.

### Serum samples

Anonymized serum samples were obtained from random blood donors in Stockholm, as previously described^4^ from week 34, 2023 (Aug. 21 - 27) and week 39, 2022 (Sep. 26 - Oct. 2), prior to the arrival of XBB in Sweden. No demographic information is available for these samples.

### SARS-CoV-2 genome analysis

The list of countries where BA.2.86 has been observed so far was obtained from metadata associated with 29 sequences available on GISAID up to Sept 1, 2023, via http://gisaid.org/EPI_SET_230901zs. The lineage growth estimates overlaid upon the phylogeny in Fig. A were performed as previously described^5^ and maintained at https://github.com/MurrellGroup/lineages (the specific model run is archived here: https://github.com/MurrellGroup/lineages/tree/b9d71cc248b61d183d1662972cfa67aa4fdeb023).

This variant competition analysis used the global GISAID SARS-CoV-2 dataset, downloaded as a bulk .fasta file on 2023-08-24, which was subsequently filtered to retain any sequences with collection dates within the previous 100 days.

The phylogeny in Fig. A is a sub-tree of the curated Nextclade^6^ BA.2 phylogeny, where leaf nodes are retained if they are among the current rapidly-growing lineages (large, coloured circles), or if they are key ancestral variants, or BA.2.86, included for context (small grey circles). The phylogeny is pruned and visualised using the MolecularEvolution.jl Julia package: https://github.com/MurrellGroup/MolecularEvolution.jl

## Ethical Statement

The blood donor samples were anonymized and therefore could be analysed without informed consent from the donors, as per advisory statement 2020–01807 from the Swedish Ethical Review Authority and the Swedish Ethics Review Act.

## Statistical analysis

Neutralising titres between groups were compared using a Wilcoxon matched-pairs signed rank test in Prism v9.

## Author contributions

Conceptualization, D.J.S., B.M.; Formal analysis, D.J.S., K.S., B.M; Conducted the assays, D.J.S., Y.Y, M.W, S.Ö., T.P.P.; Designed the methodology, D.J.S., K.S., T.P.P, B.M.; Responsible for figures and tables, D.J.S., T.P.P., B.M.; Resources, S.M., G.B.K.H., J.A., B.M.; Oversaw the study, D.J.S., B.M; Funding Acquisition, G.B.K.H., J.A., T.P.P., B.M. Writing – original draft, D.J.S., B.M.; Writing – review & editing, D.J.S., T.P.P., G.B.K.H., J.A., B.M. D.J.S and B.M. were responsible for the decision to submit the manuscript for publication.

